# Factors affecting the survival of tree seedlings following shifting cultivation in the Eastern Himalaya

**DOI:** 10.1101/619981

**Authors:** Karthik Teegalapalli, Rohan Arthur, Suhel Quader, Aparajita Datta

## Abstract

1. The seedling stage is considered an important bottleneck determining forest community composition during succession. Seedling growth and survival are known to be affected by factors such as light availability, herbivory and competition with existing vegetation. A shifting cultivation landscape with a cycle of clearing and regeneration provides an experimental setup to understand secondary succession.
2. We introduced tree seedlings of three species with differing life history strategies in an uncut forest and in sites formed 2, 6, 12, 25, 50 and 100 years following shifting cultivation in a subtropical landscape in the Eastern Himalaya. Seedlings of a pioneer species *Saurauia nepalensis*, a mid-successional species *Terminalia myriocarpa* and a mature forest species *Castanopsis indica* were introduced in plots with a control and three treatments: shading (light manipulated), clearing of vegetation (reduced competition), and exclosures (mammal herbivory excluded) and monitored for 18, 22 and 23 months, respectively.
3. *Saurauia* survived relatively well in all sites, *Terminalia* seedlings survived relatively well in the 12 and 25 year sites, and *Castanopsis* seedlings survived well in the 100 year site and the uncut forest. In the early successional site, survival of *Saurauia* was higher within exclosures indicating negative effects of herbivory. In the same site, clearing vegetation improved the survival of *Terminalia*, implying competition with existing vegetation. *Castanopsis* had the highest survival in uncut forest and survival was almost nil in the early successional site.
4. Synthesis: Seedling survival in a successional landscape depended on species-as well as site-specific factors. The existing vegetation in an early successional site had a negative effect on the survival of the mid-successional species, while herbivory had a negative effect on the pioneer species. Survival of the mature forest species in the early successional site was negligible but was high in old successional and uncut forest sites, indicating that mature forest species can colonize sites and survive only after certain physical and biotic aspects of the environment have been met. Our experiments were useful to examine factors that affect survival of tree species with differing life history strategies in a shifting cultivation landscape.

## Introduction

Secondary succession of plants has interested biologists for over a century since Clements (1916) and Gleason (1917) suggested their contrasting theories: one, a holistic theory with a climax equilibrium stage and the other an individualistic one with an open-ended view of succession. Egler (1954) suggested that vegetation development occurs in one of two ways: relay floristics, in which one species establishment relays to the next and initial floristics, in which different species establish in a site and assume predominance at different stages of development. In terms of the processes during succession, Connell and Slatyer (1977) suggested three distinct mechanisms of species interactions: facilitation, tolerance and inhibition, while Pickett, Collins and Armesto (1987) suggested a combination of these at different successional stages. More recent research has focussed on additional factors that affect successional patterns such as: 1) the disturbance regime; the scale and frequency of disturbances (Denslow, 1980; Lawrence et al., 2005), 2) environmental factors such as soil and climate (Bazzaz, 1979; Pickett, Collins & Armesto, 1987), 3) biotic factors such as herbivory (Coley, 1983), phylogenetic structure of plant communities (Letcher et al. 2014), life history strategies (Bazzaz, 1979; Denslow, 1980) and functional traits of species (Schleicher, Peppler-Lisbach & Kleyer 2011).

These factors can be expected to be species- and site-specific in different seral stages of succession (Bazzaz, 1979; Uhl, 1987). Initially, during succession from a clearing to a forest, early successional species may establish and survive since they are better adapted to the harsh and variable environmental conditions, whereas propagules of mature forest species and old successional species may fail to germinate and survive in such sites (Bazzaz, 1979). Among the environmental factors that affect species differentially, light availability has been shown to limit tree seedling recruitment of some species while facilitating that of others (Beckage, Lavine & Clark, 2005; Walters & Reich, 1996). Herbivory by vertebrate browsers and invertebrates, which is considered an important bottleneck in the regeneration stage, can also be expected to vary in different successional stages (De Steven, 1991; Coley & Barone, 1996). Insect herbivory has been shown to be high during the early years of succession, with herbivory assemblages changing from foliar-feeding insects in early stages to sap-feeding insects in the later stages of succession (Brown & Gange, 1992, Carson & Root, 1999). While mammal herbivory can be higher in early successional sites due to higher visibility of seedlings, it can also be higher in older successional sites and forests where the vertebrate abundance is usually higher (Tabarelli & Peres, 2002). Existing vegetation in an early successional site may foster survival of pioneer species while hindering that of mature forest species (Davis, Wrage & Reich, 1998; Putz & Canham, 1992).

These theories pertaining to succession mechanisms and the role of biotic and abiotic factors can be tested by introducing propagules of species with different life history strategies in mature forest and different-aged successional sites, manipulating different factors in plots within sites and monitoring their survival over time. Several studies in the past have undertaken such experiments (*e.g*. Armesto & Pickett, 1985; Beckage & Clark, 2003; Berkowitz, Charles & Victoria, 1995; De Steven, 1991; Uhl, 1987), however field experiments from the Old World are few (Fayolle et al., 2015, Goodale et al., 2014; Khan & Tripathi,1989, Lin et al., 2014). These multi-factor experiments are difficult to simulate in natural conditions. However, a shifting cultivation landscape, which involves a continuous cycle of clearing and secondary succession, creating a landscape where spatially separated sites can represent temporal stages of succession, provides such a quasi-experimental setup. The succession phase, however, is relatively short due to repeated cycles of cultivation and site manipulation for cultivation can alter natural succession patterns. Despite these shortcomings, studying succession at such sites has practical advantages: while patterns in older sites can be used to infer succession over relatively long periods of time, successional sites representing a range of different-aged sites can be used to identify temporal trends in succession (Pickett, 1989).

We used an artificially maintained seral stage gradient (from recently cleared to mature forest) to measure seedling success of three tree species with contrasting life history traits. We chose a pioneer, an intermediate and a mature forest species based on the relative abundances of the species in different successional sites and uncut forest in the study site (data from Teegalapalli & Datta 2016). We undertook a series of controlled experiments to examine the effects of light availability, herbivory and competition on survival across this gradient. The specific objectives of our research were: 1) to understand the effects of the age of a successional site on the seedling survival of a mature, a pioneer and a mid-successional tree species, and 2) to understand the effects of shade, existing vegetation and mammalian herbivory on the seedling survival of these species in different-aged successional sites and uncut forest. Based on existing literature, we predicted that: 1) the three species will have higher seedling survival in the successional sites where the adults of the species were relatively abundant, 2) herbivory by mammals will negatively affect seedling survival in the early successional site, 3) providing artificial shade will improve seedling survival, particularly for the mature species in the early successional sites, and, 4) clearing aboveground vegetation in the early successional sites will improve seedling survival of the three species by reducing competition.

## Materials and methods

### Study site

We undertook our experiments in a shifting cultivation landscape around Bomdo village in the Eastern Himalaya (Figure 1). The village is located in the Upper Siang district of the state of Arunachal Pradesh (India) at 28.753° N and 94.896° E. The region receives relatively high rainfall of over 4000 mm every year (Anonymous, 2015) and features tropical wet- and semi-evergreen forests and sub-tropical broad-leaved forests in the areas above 800 msl (Singh et al., 1996). The resident *Adi* community practice shifting cultivation in a rotational system in three large blocks around the village. The fallow period, the gap between two cultivation cycles is at least 10 years, as shown in other sites in the Upper Siang district (Borang, 1997). Some of the patches in the landscape have been abandoned for relatively long periods (over 100 years) due to factors such as inaccessibility and low productivity. The *Adi* maintain a systematic oral cultivation history based on which we selected successional sites that were cultivated and left fallow for 2, 6, 12, 25, 50 and about 100 years and an uncut forest site (hereafter, referred to as uncut forest) in the landscape for comparison. Uncut forest in this study represents moderately disturbed uncut forest rather than a mature forest that is devoid of any human disturbance. Hereafter, we refer to the 2 and 6 year sites as early, the 12 and 25 year sites as intermediate and the 50 and 100 year old sites as old successional sites. The sites were located within 1000 m distance of each other and ranged in elevation from 750 to 1100 msl. Prior permission to undertake the research in the shifting cultivation landscape around the village was sought from the village elders.

**Figure 1:**
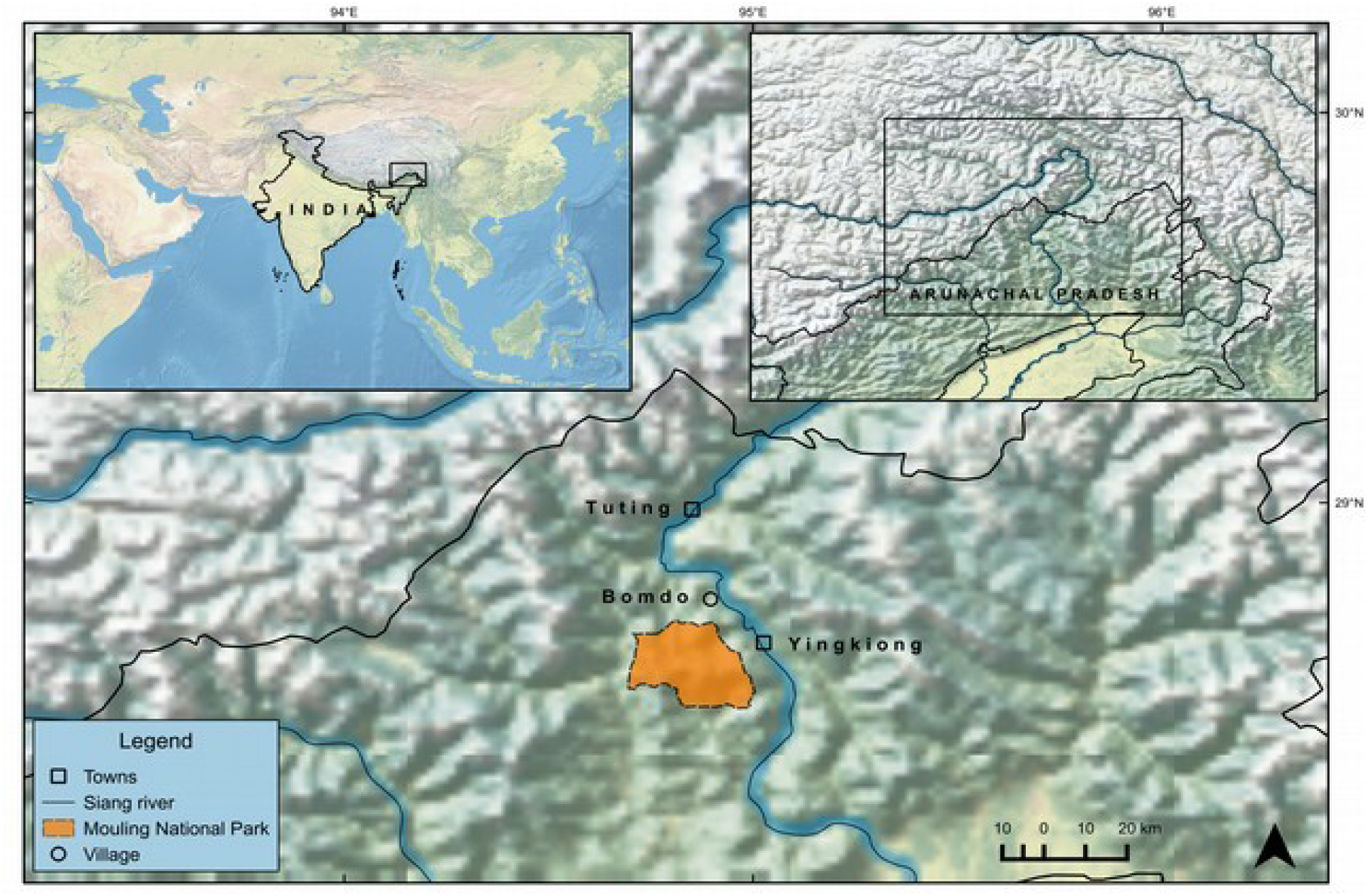
Map depicting the location of the study area in the Arunachal Pradesh state in north-east India. The manipulative experiments to understand factors that affect tree seedling survival were undertaken in the shifting cultivation landscape around the Bomdo village.

### Study species

We used three species with differing life-history strategies: *Castanopsis indica* (Roxb.) A. DC. (Family Fagaceae), *Saurauia nepalensis* Wall. (Family Actinidiaceae) and *Terminalia myriocarpa* Van Huerck et Muell.-Arg. (Family Combretaceae) for the experiments. These species were selected based on vegetation data collected from 2, 12, 25 and 50 year sites and uncut forest, for another study (Teegalapalli & Datta, 2016). *C. indica*, (hereafter, *Castanopsis*) is a shade-tolerant mature forest species that was recorded only from the uncut forest and comprised about 6 % of all individuals in that forest. *C. indica* and two more species from the genus *Castanopsis* comprised about a fourth of all individuals recorded from the uncut forest. *S. nepalensis*, (hereafter, *Saurauia*), a light-loving pioneer species was a dominant species in the early successional sites comprising 70 % and 44 % of all trees recorded from 6 and 12 year old sites, and was not recorded from the 100 year site and uncut forest. *T. myriocarpa*, (hereafter, *Terminalia*), a fast-growing hardwood species (Bhattacharya & Nanda, 2005; Qureshi, 1968) was recorded in low numbers, only from intermediate successional sites (12 and 25 year sites), and is considered as a mid-successional species. While in this site it was only recorded from intermediate successional stages, the species has been recorded as a dominant tree in tropical semi-deciduous forests in Arunachal Pradesh and other parts of North-east India (Bhuyan, Khan & Tripathi et al., 2003).

### Experimental setup

We used a multifactorial design with three treatments within each block: clearing of above-ground vegetation to reduce competition with existing vegetation, artificial shade to manipulate available light and exclusion of mammal herbivory across different-aged sites and uncut forest. Although insect herbivory is an important factor affecting seedling survival in tropical forests (Coley & Barone, 1996; Crawley, 1983), we did not test for this factor because we refrained from spraying chemicals to exclude insect herbivory in a shifting cultivation site that was actively used for agriculture by the local community. We used a block design with eight 1 × 1 m^2^ plots (seven plots for three treatments, singly and in combination, and a control plot) within each 2 × 4 m^2^ block (Figure 2). For *Castanopsis* and *Saurauia*, we had three replicates of the blocks and for *Terminalia* we had five replicates each in the 2, 6, 12, 25, 50, 100 year sites and the uncut forest site.

**Figure 2:**
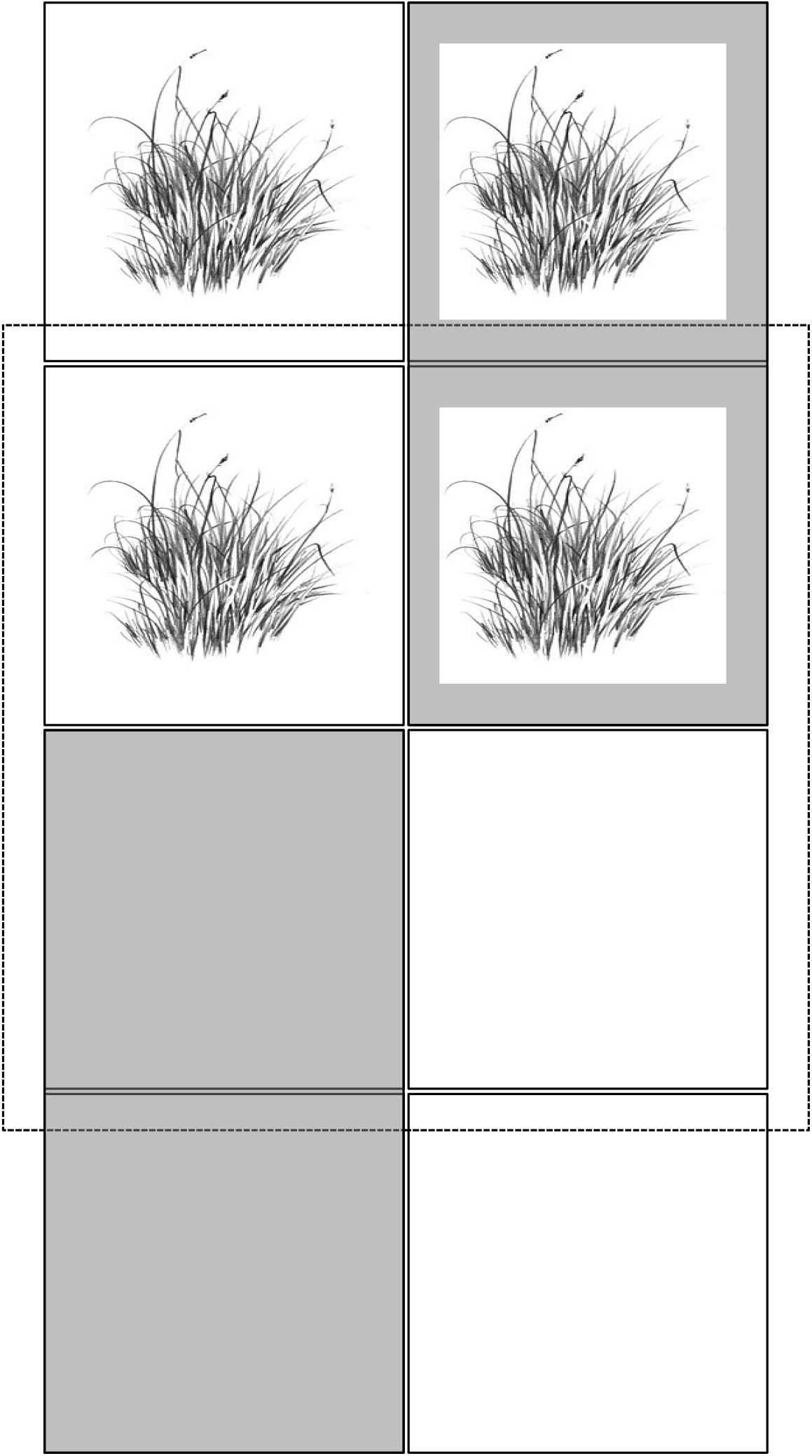
The multi-factorial block design depicting shaded plots (grey), vegetation cleared plots in which above-ground vegetation was cleared quarterly (bottom four plots) and plots with exclosures (dotted lines). Seedlings were introduced in each of the 1 × 1 m plots nested within the 2 × 4 block.

In the treatment plots, we controlled for mammal herbivory by fencing the plot using metal wire-mesh exclosures. The primary mammalian herbivores in the study area are barking deer (*Muntiacus muntjak*) and wild pig (*Sus scrofa*). Apart from wild mammalian herbivores present in the landscape, the *Adi* community rears domestic pigs and a species of bovid locally called *Mithun (Bos frontalis*), which are free-ranging livestock. We manipulated available light using 40 % artificial greenhouse shade-netting (leaf litter collected in the net was cleared monthly) and regularly cleared above-ground vegetation (once every three months) to eliminate competition with above-ground vegetation.

First year seedlings (5–10 cm height) of the three species chosen for the experiments were collected from regenerating secondary and uncut forests in the landscape and planted in the growing season (between March and June) in the different-aged successional sites and uncut forest. We avoided disturbing the roots of the seedlings to prevent root-shock and planted the seedlings within two hours of collection from secondary forests. Overall, for *Castanopsis* and *Saurauia*, we transplanted 5 seedlings per 1 × 1 m^2^ plot, 40 seedlings in each block, 120 seedlings in each successional site and the uncut forest, a total of 840 seedlings each across the study site. For *Terminalia*, which had five replicates in each site, we transplanted a total of 1400 seedlings. Based on the availability of the different species of seedlings from natural regeneration, *Castanopsis* seedlings were transplanted in April 2011, *Saurauia* seedlings were transplanted in May 2011 and seedlings were monitored till February 2013. *Terminalia* seedlings were transplanted in July 2012 and monitored till November 2013. Seedling survival was monitored monthly and recorded as alive or dead for a period of 23, 22 and 18 months for *Castanopsis, Saurauia and Terminalia*, respectively.

### Analytical methods

#### Across all sites

Generalized Linear Mixed Models (GLMMs) were used with the proportion of seedlings alive at the end of the study in each 1 × 1 m plot as the response variable (varying from 0 to 1). Fixed effects were of the different treatments (singly and in combination) and age of site, and within-site block identity was included as a random effect. To use ‘age of the site’ as a continuous variable for the analysis, the uncut forest was assigned an age of 300 years (*e.g*. Raman, Rawat & Johnsingh, 1998; Riswan, Kenworthy & Kartawinata, 1985).

#### Within a site

We used GLMMs with the seedling survival status of each seedling at the end of the study (dead or alive) as the response variable, the different treatments, singly and in combination as fixed effects and variation between blocks as well as variation between seedling replicates as random effects to investigate the importance of treatments within different-aged sites and uncut forest. We used the statistical software R for the analyses with the ‘survival’ package for survival analysis (Therneau, 2015) and ‘lme4’ package (Bates et al., 2015) for GLMMs (R Development Core Team, 2015).

#### Survival curves

We used Kaplan-Meier (K-M) survival curves for plotting seedling survival patterns (Kaplan & Meier, 1958). The curves were used to visualise the right-censored survival data (some of the seedlings were still alive at the end of the study) of seedlings in control and treatment plots and log rank tests were used to determine if the curves differed statistically, which indicated the effects of different treatments on the survival period of seedlings. Log rank tests are similar to the Chi-square tests, in which a test statistic is calculated to test the null hypothesis that the survival curves are similar for the groups being compared.

## Results

### General patterns

Across all sites, overall seedling mortality was 7 %, 23 % and 35 % within the first month for *Saurauia, Castanopsis* and *Terminalia*, respectively, which increased to 25 %, 30 % and 55 % in the first three months. A majority of the mortality in the first quarter was from the early successional sites. About a third of the seedlings of *Castanopsis* and *Saurauia* and about a fourth of the seedlings of *Terminalia* survived till the end of the study. While few *Saurauia* seedlings survived in the control plots in the 2 year site (13 % survival), seedlings in the control plots survived relatively well in the 6 (40 % survival) and 12 year (33 % survival) sites as well as in the 100 year site (47 %) and the uncut forest (33 %, Figure 3). The survival of *Castanopsis* was very low in the control plots in the early successional sites (nil in the 2 year site and 13 % survival in the 6 year old site) and more than 50 % in the 100 year site and the uncut forest (Figure 3). About a fourth of *Terminalia* seedlings survived in the control plots in the 2 year site, the 6 year site and uncut forest while survival was highest in the 12 year site (56 %, Figure 3).

**Figure 3:**
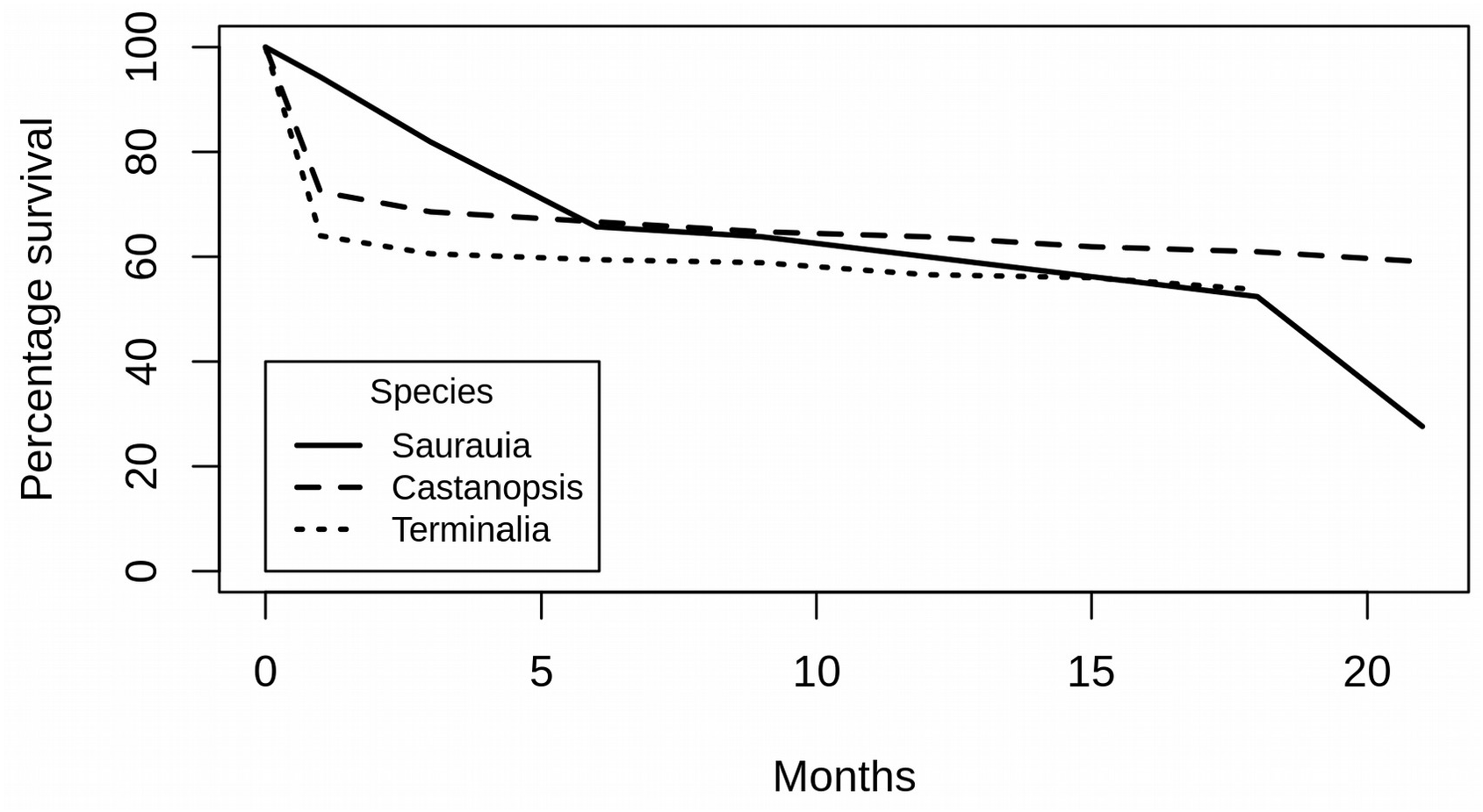
Overall survival of *Saurauia nepalensis* (N=105), *Castanopsis indica* (N=105) seedlings over a period of 21 months and *Terminalia myriocarpa* (N=175) seedlings in the control plots over a period of 18 months, averaged across different ages.

### Seedling survival across all sites

The output of the GLMM undertaken across sites with the age of site and different treatments as predictor variables and proportion of seedlings alive at the end of the study as the response variable indicated that while the age of site did not have a significant effect on *Saurauia* and *Terminalia* seedling survival, survival of *Castanopsis* increased with age (*F* = 0.01, *p* = 0.005, Figure 4, Table 1, full model results in Table S1 – S3). Across all sites, survival of *Saurauia* was higher in exclosure plots (*F* = 1.83, *p* = 0.01, Table 1). None of the treatments had a significant effect on the survival of *Terminalia*.

**Figure 4:**
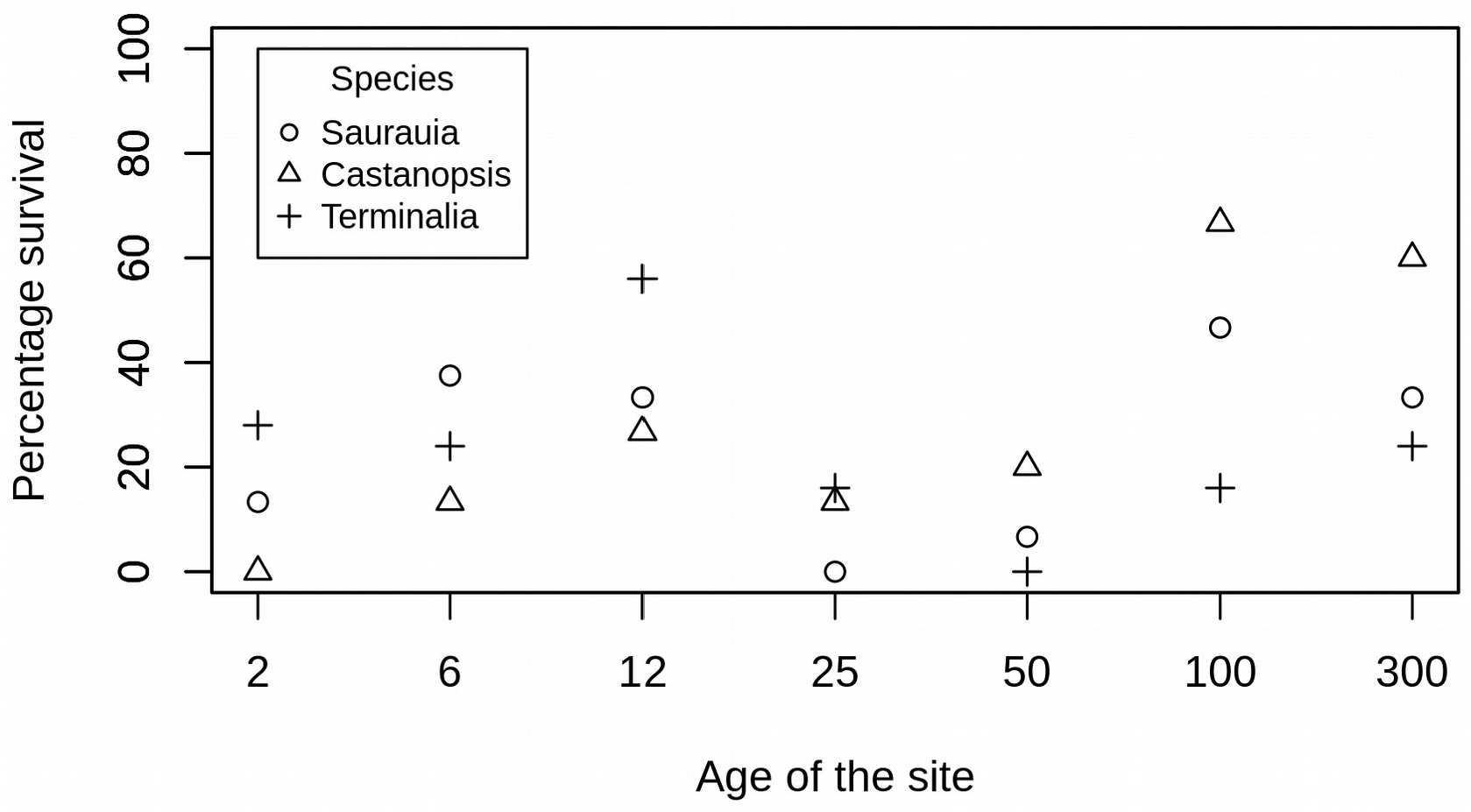
Survival of *Saurauia nepalensis* (n=105), *Castanopsis indica* (n=105) and *Terminalia myriocarpa* (n=175) in the control plots in successional sites formed following 2, 6, 12, 25, 50 and 100 years following shifting cultivation. The uncut forest site has been assigned an age of 300 years to show the survival of the three species in the site.

**Table 1:** Generalised Linear Mixed Models results for seedling survival of *Castanopsis indica, Saurauia nepalensis and Terminalia myriocarpa* seedlings (pooled across each 1 × 1 m plot) at the end of the study across successional sites formed 2, 6, 12, 25, 50 and 100 years following shifting cultivation and uncut forest and treatments (* indicates a *p*-value < 0.05). Proportion of seedlings alive at the end of the study was the response variable. The factors examined were shade, herbivory and competition with aboveground vegetation. Greenhouse shade netting was used to manipulate available light, exclosures were used to prevent herbivory by mammals and vegetation was cleared to eliminate competition with existing vegetation. Parameter estimates (intercept and contrast), standard errors (SE) and hypothesis tests for parameters are shown.

| *Castanopsis indica* across all sites and treatments |  |  |  |  |
| --- | --- | --- | --- | --- |
|  | Estimate | S.E. | z value | *p* |
| Age | 0.01 | 0.004 | 2.8 | 0.005** |
| *Saurauia nepalensis* across all sites and treatments |  |  |  |  |
| Exclosures - herbivory | 1.81 | 0.72 | 2.52 | 0.01** |
| *Terminalia myriocarpa* across all sites and treatments |  |  |  |  |
| No effects of age or treatment on seedling survival |  |  |  |  |

### Seedling survival within sites

GLMM analysis undertaken in different-aged sites and uncut forest indicated factors that affected seedling survival specifically in each site. For *Saurauia* in the 2 year site, exclosure plots singly (*F* = 2.39, *p* = 0.02) and in combination with shade (*F* = 2.10, *p* = 0.03) had significantly higher seedling survival in comparison with control plots (Table 2). Exclosure plots in uncut forest (*F* = 2.91, *p* = 0.00) also had higher seedling survival than control plots (Table 2).

**Table 2:** Generalised Linear Mixed Models results for seedling survival of *Saurauia nepalensis* seedlings (pooled across each 1 × 1 m plot) at the end of the study within successional sites formed 2, 6, 12, 25, 50 and 100 years following shifting cultivation and uncut forest (only sites where specific factors had a significant effect on seedling survival have been shown, * indicates a *p*-value < 0.05). The factors examined were shade, herbivory and competition with aboveground vegetation. Greenhouse shade netting was used to manipulate available light, exclosures were used to prevent herbivory by mammals and vegetation was cleared to eliminate competition with existing vegetation. Parameter estimates (intercept and contrast), standard errors (SE) and hypothesis tests for parameters are shown.

| ***Saurauia nepalensis* in 2 year site** |  |  |  |  |
| --- | --- | --- | --- | --- |
| | Estimate | S.E. | z value | $p$ |
| Intercept | -1.96 | 0.83 | -2.35 | 0.02* |
| Exclosures | 2.39 | 1.04 | 2.29 | 0.02* |
| Shade + Exclosures | 2.10 | 1.04 | 2.03 | 0.04* |
| ***Saurauia nepalensis* in uncut forest site** |  |  |  |  |
| Exclosures | 2.91 | 1.00 | 2.90 | 0.004* |

For *Terminalia*, vegetation cleared plots (*F* = 1.42, *p* = 0.04) and plots with all three treatments (*F* = 1.78, *p* = 0.01) had higher seedling survival than control plots in the 2 year site (Table 3). For *Terminalia*, there were unexpected results with the treatments in the 12 year site; shade and vegetation cleared plots had lower survival than control plots (*F* = −1.54, *p* = 0.02, Table 3).

**Table 3:** Generalised Linear Mixed Models results for survival of *Terminalia myriocarpa* seedlings (pooled across each 1 × 1 m plot) at the end of the study within successional sites formed 2, 6, 12, 25, 50 and 100 years following shifting cultivation and uncut forest (only sites where specific factors had a significant effect on seedling survival have been shown, * indicates a *p*-value < 0.05). The factors examined were shade, herbivory and competition with aboveground vegetation. Greenhouse shade netting was used to manipulate available light, exclosures were used to prevent herbivory by mammals and vegetation was cleared to eliminate competition with existing vegetation. Parameter estimates (intercept and contrast), standard errors (SE) and hypothesis tests for parameters are shown.

| ***Terminalia myriocarpa* in 2 year site** |  |  |  |  |
| --- | --- | --- | --- | --- |
|  | Estimate | S.E. | z value | *p* |
| Intercept | -0.99 | 0.50 | -1.97 | 0.05* |
| Vegetation cleared | 1.42 | 0.69 | 2.06 | 0.04* |
| Shade + Exclosures +<br>Vegetation cleared | 1.78 | 0.70 | 2.53 | 0.01* |
| ***Terminalia myriocarpa* in 12 year site** |  |  |  |  |
| Shade + Vegetation<br>cleared | -1.54 | 0.65 | -2.37 | 0.02* |

For *Castanopsis*, in the 25 year site, shade, singly (*F* = 3.00, *p* = 0.00) and in combination with exclosures (*F* = 2.23, *p* = 0.03) as well as in combination with all treatments (*F* = 3.85, *p* = 0.00) had higher survival than control plots (Table 4).

**Table 4:** Generalised Linear Mixed Models results for survival of *Castanopsis indica* seedlings (pooled across each 1 × 1 m plot) at the end of the study within successional sites formed 2, 6, 12, 25, 50 and 100 years following shifting cultivation and uncut forest (only sites where specific factors had a significant effect on seedling survival have been shown, * indicates a *p*-value < 0.05). The factors examined were shade, herbivory and competition with aboveground vegetation. Greenhouse shade netting was used to manipulate available light, exclosures were used to prevent herbivory by mammals and vegetation was cleared to eliminate competition with existing vegetation. Parameter estimates (intercept and contrast), standard errors (SE) and hypothesis tests for parameters are shown.

| ***Castanopsis indica* in 25 year site** |  |  |  |  |
| --- | --- | --- | --- | --- |
|  | Estimate | S.E. | z value | *p* |
| Intercept | -2.45 | 1.15 | -2.12 | 0.03* |
| Shade | 3.00 | 1.06 | 2.82 | 0.00* |
| Shade + Exclosures | 2.23 | 1.03 | 2.15 | 0.03* |
| Shade + Exclosures +<br>Vegetation cleared | 3.85 | 1.12 | 3.43 | 0.00* |

### Survival curves

The Kaplan-Meier (K-M) survival curves indicated species-specific as well as treatment-specific differences, which were tested using the log-rank tests, based on which statistically significant differences (*p* < 0.05) between seedling survival in control and treatments plots were identified (Figures S1 – S3, Tables S4 – S6). All other survival curves that were not significant have not been shown.

For the pioneer species *Saurauia*, survival curves in the herbivory exclosure plots in the 2 and 25 year sites and uncut forest were significantly different from those in control plots, respectively (Figure S1). Seedling survival curves of plots with all three treatments combined were significantly different in the 2 year and 50 year site than those in control plots for *Saurauia*. Survival curves in vegetation cleared and control plots the 6 year site were significantly different. Further, survival curves in treatments with shade and vegetation clearing were different from those in control plots in the 12 year and 100 year sites. In the 50 year site, survival curves were significantly different in exclosure plots with vegetation clearing and shade plots in comparison with those in control plots.

For *Terminalia*, in the 2 year site, survival curves in vegetation cleared plots and plots with all three treatments combined were significantly different from those in control plots (Figure S2). In the 12 year site, the survival curves in the shade and vegetation cleared plots were significantly different from those in control plots. In the 25 year sites and the older successional sites and uncut forest, the survival of seedlings in control and treatments plots was lower than 20 % and survival curves for these sites were not plotted.

For *Castanopsis*, in the 6 year site, survival curves were significantly different between control plots and plots with exclosures and vegetation cleared and plots with all three treatments (Figure S3). In the 25 year site, survival curves were significantly different between control plots and plots with shade and plots with all three treatments. In the 100 year site, plots in which vegetation was cleared had significantly different survival curves in comparison with control plots.

## Discussion

### Seedling survival patterns

Overall, more than a quarter of the seedlings introduced in this study survived till the end of the study and seedling mortality was highest in the first six months, with the highest mortality in the early successional site. Seedlings of the pioneer species *Saurauia* survived generally well in all sites (20 – 35 %) except the 25 year site (~8 %), the mid-successional species *Terminalia* seedlings survived relatively well in the 2 year site (~45 %) and the two intermediate sites (6 year [~39%] and 12 year [~47%] sites), and the mature forest species *Castanopsis* seedlings survived well in the 100 year (~53 %) site and the uncut forest (~66 %).

Several *Castanopsis* species are characteristic of mature sub-tropical forests in the Eastern Himalaya (Sundriyal et al., 1994), therefore we expected the survival of the species to be highest in the uncut forest. However, we did not expect that *Saurauia* seedlings would survive well in the older sites and uncut forest, given that it is known to be a pioneer species. Although *Saurauia* seedlings survived in these sites till the end of the study, it is unlikely that they will survive to the adult stage, since we recorded trees of the species in 12 year and 25 year successional sites and did not find trees or seedlings of the species in the vegetation sampling we undertook in the older successional sites and uncut forest (Teegalapalli & Datta, 2016). Similarly, *Saurauia* trees were recorded from early successional fallows formed following shifting cultivation in Vietnam but were not recorded from adjoining old-growth evergreen broad-leaved forest (Van Do, Osawa & Thang, 2010). We speculate that other mortality factors at later stages (sapling or pole) that we did not monitor in our study may result in the absence of this species in late successional sites. The seedling survival patterns of *Terminalia* reflected the adult tree composition patterns recorded from the site: trees of the species were only recorded from the intermediate (6 and 12 year sites) successional sites (Teegalapalli & Datta, 2016).

### Factors affecting seedling survival

The pioneer species *Saurauia* was more susceptible to herbivory: introducing exclosures that prevented herbivory by mammals improved the number of seedlings that survived as well as the survival period across all sites. The ‘strategic resource allocation’ concept (McCook, 1994) suggests that pioneer species expend more on faster growth than on herbivory defense in contrast with slow-growing mature forest species that need to defend themselves due to their slower growth and expend more on secondary metabolites (Coley, 1983; Feeny, 1976). The higher rates of growth have been attributed to better adaptation to nutrient intake and retention under low soil nutrient conditions and allocation of high amounts of energy to root production (Uhl, 1987). For example, in Barro Colorado Island, Panama, pioneer species had six times higher levels of herbivory than mature forest persistent species (Coley, 1983). In this study site, herbivory is likely an important factor affecting regeneration since the livestock that the resident *Adi* community rear graze in the successional fallows*. Saurauia* can be expected to be particularly susceptible to herbivory as the species has relatively high calorific value and is also used as cattle fodder in Sikkim and Nepal (Chettri & Sharma, 2008).

Survival of the mature forest species *Castanopsis* was enhanced in plots with all three treatments combined; shade, exclosures and vegetation cleared in the 6 and 25 year sites. Specifically, providing shade in the 25 year old site improved the survival of the species, while seedlings in the 6 and 12 year sites had relatively low survival both in control and shade plots. This indicates that the favourable conditions required for this shade-loving species to survive were not met even in shaded plots that provided 40 % of the ambient light conditions in the intermediate successional sites. However, *Castanopsis* seedlings surviving better in vegetation cleared plots than in control plots could be due to factors that were not controlled for in this study.

*Terminalia* seedlings survived relatively well in the intermediate sites. *T. myriocarpa* is a fast-growing, light-tolerant species (Bhattacharya & Nanda, 2005; Deb et al., 2014; Qureshi, 1968) and relatively less affected by herbivory as shown in this study, which likely fosters its survival in the intermediate successional sites. In the 2 year old site, however, both clearing vegetation, singly as well as in combination with shade and herbivory exclosures improved survival of the seedlings, indicating that the species is unable to colonize recently abandoned clearing due to a combination of factors.

### Insights into succession mechanisms

Pickett, Collins and Armesto (1987) suggested that a combination of mechanisms can be applicable in a successional stage in contrast with the individualistic models of facilitation, tolerance and inhibition suggested by Connell and Slatyer (1977). In our study, seedling survival patterns in different treatments indicated species-specific as well as site-specific interactions between species and the existing vegetation. Introducing exclosures and shade positively affected seedling survival of *Saurauia* in the early successional site indicating that pioneer tree species cannot immediately colonize an open area due to the harsh physical conditions and may need to be facilitated by the pioneer shrubs that colonize the clearings first.

The vegetation in the 2 year site was dominated by seedlings and saplings of the small tree / shrub species *Maesa indica* and *M. ramentaceae* (Teegalapalli & Datta, 2016). Fruits of these species are small and widely dispersed by smaller frugivorous birds and *Maesa indica is* a forest-gap adapted early-successional species in southern India (Chetana & Ganesh, 2012; Ganesh & Davidar, 2001). These species possibly modify the environment to make it suitable for the early successional tree species to colonize, which indicates facilitation. A similar successional trend was recorded in abandoned fields in the Rio Negro region of the Amazon Basin (Uhl, 1987); grasses and forbs dominated in the first year of succession, followed by a dominance of shrubs and long-lived tree species were recorded only after 5 years.

In contrast, clearing existing vegetation in the early successional site improved the survival of *T. myriocarpa* suggesting competition as the likely mechanism inhibiting its survival. However, it was difficult to clearly tease apart if competition with existing vegetation was the single factor that improved survival of seedlings since seedlings in plots with all three treatments also survived better than control plots in the early successional site. This is particularly so, since in the 12 year site, contrary to expectations, the combined effect of providing shade and clearing vegetation reduced the survival of the species. Survival of the mature forest species *Castanopsis* seedlings in control and treatment plots in the early successional site was negligible, indicating that mature forest species can colonize sites, survive and reach the adult stage only after certain conditions of the physical environment have been met.

## Site-specific factors that can affect seedling survival

A shifting cultivation landscape such as the one used for this research is one that is in a constant flux of farming management. While the early successional sites are sites that were recently cultivated, the intermediate sites are the ones that will be cultivated in the future since the fallow period, the gap between two cultivation phases, is 10 years in the region (Teegalapalli & Datta 2016, Teegalapalli et al. 2018). However, the older 25 and 50 year sites are sites which were cultivated once and abandoned subsequently due to low soil fertility and crop productivity. Therefore, it is likely that seedling survival in these sites do not reflect the seedling survival in a site that has recovered for 25 to 50 years. On the contrary, the 100 year old site in the landscape chosen for this research was one which was not abandoned for the same reasons and seedling survival in this site was comparable with that in uncut forest.

## Conclusion

To our knowledge, this is the first study in India in which manipulative experiments were undertaken to investigate the importance of multiple factors that affect seedling survival in a successional landscape. The mature forest tree species did not survive in the early successional sites, whereas under certain conditions, the mid-successional and the pioneer species survived relatively well. The seedling survival of the mature species increased with age and was highest in uncut forest. Mammalian herbivory was an important factor affecting the pioneer species. Existing vegetation negatively affected the survival of *Terminalia* seedlings in the early successional site indicating competition with existing vegetation and more than half of the seedlings of the species survived in the intermediate successional sites from where adult trees of the species were also recorded. This study provided insights into understanding the relative effects of light availability, herbivory and existing vegetation on the seedling survival of a mature forest, a mid-successional and an early successional tree species in successional sites and uncut forest.

## Acknowledgments

The ATREE Small Grants for Research in North-east India, Ravi Sankaran Inlaks Foundation, Idea Wild and the Rufford Foundation funded the research presented. The first author was supported by the DBT-RA program in Biotechnology and Life Sciences during the writing of the manuscript. Dunge Yalik, Army Duggong, Gekut Medo and Bamut Medo helped with the intensive fieldwork involved in the research. The Arunachal Pradesh Forest Department: Shri Bittem Darang (Divisional Forest Officer) and Shri Kopang Takuk (Forest Ranger) provided logistical help. T R Shankar Raman provided useful comments to the manuscript. We thank these individuals and organisations for their support.

## Authors’ contributions

KT, RA, SQ and AD conceived the idea and designed the methodology. KT collected the data in the field with inputs from AD, analysed the data with inputs from SQ and wrote the manuscript with inputs from RA, AD and SQ. All authors contributed to the drafts and gave final approval for publication.

## Data accessibility

All the data used in the research presented in the manuscript will be archived in Data Dryad.

## References

Anonymous. (2015). Indian Meteorological Department. www.imd.gov.in, accessed on 20th August, 2015.

Armesto, J. J. & Pickett, S. T. A. (1985). Experiments on disturbance in old-field plant communities: impact on species richness and abundance. Ecology, 66, 230–240.

Bates, D., Maechler, M., Bolker, B. & Walker, S. (2015). Fitting Linear Mixed-Effects Models Using lme4. Journal of Statistical Software, 67, 1–48. doi: 10.18637/jss.v067.i01.

Bazzaz, F. A. (1979). The physiological ecology of plant succession. Annual Review of Ecology and Systematics, 10, 351–371.

Beckage, B., Lavine, M. & Clark, J. S. (2005). Survival of tree seedlings across space and time: estimates from long-term count data. Journal of Ecology, 93, 1177–1184. doi: 10.1111/j.1365-2745.2005.01053.x

Beckage, B. & Clark, J. S. (2003). Seedling survival and growth of three forest tree species: the role of spatial heterogeneity. Ecology, 84 (7), 1849–1861. doi: https://doi.org/10.1890/0012-9658(2003)084[1849:SSAGOT]2.0.CO;2

Berkowitz, A.R., Charles D. C., & Victoria R. K. (1995). Competition vs. facilitation of tree seedling growth and survival in early successional communities. Ecology, 76, 1156–1168. doi: https://doi.org/10.2307/1940923

Bhattacharya, B. & Nanda, S. K. (2005). Shifting cultivation in north-east India: technological alternatives and extension implications. In: Sustainable agriculture: issues in production, management, agronomy and ICT applications (Eds. Bandhopadhyay, A., Sundaram, K. V., Moni, M., Kundu, P. S. and Jha, M. M.), Northern Book Centre, New Delhi, India, 327 pages.

Bhuyan, P., Khan, M. L. & Tripathi, R. S. (2003). Tree diversity and population structure in undisturbed and human-impacted stands of tropical wet evergreen forest in Arunachal Pradesh, Eastern Himalayas, India. Biodiversity and Conservation 12, 1753–1773.

Borang, A. (1997). Shifting cultivation among the Adi tribes of Arunachal Pradesh. Journal of Human Ecology, 6, 145–151.

Brown, V. K. & Gange, A. C. (1992). Secondary plant succession: how is it modified by insect herbivory. Vegetatio, 101, 3–13. doi: https://doi.org/10.1007/BF00031910

Carson, W. P. & Root, R. B. (1999). Top-down effects of insect herbivores during early succession: influence on biomass and plant dominance. Oecologia, 121, 260–272. doi: https://doi.org/10.1007/s004420050928

Chetana, H. C. & Ganesh, T. (2012). Importance of shade trees (Grevillea robusta) in the dispersal of forest tree species in managed tea plantations of southern Western Ghats, India. Journal of Tropical Ecology, 28, 187–197. doi: 10.1017/S0266467411000721

Chettri, N. & Sharma, E. (2008). Traditional knowledge on firewood and fodder values corresponds to scientific assessment. IUFRO World Series Vol. 21, p. 31.

Clements, F. E. (1916). Plant succession: an analysis of the development of vegetation (No. 242). Carnegie Institution of Washington, DC.

Coley, P. D. & Barone, J. A. (1996). Herbivory and plant defenses in tropical forests. Annual Review of Ecology and Systematics, 27, 305–335. doi: https://doi.org/10.1146/annurev.ecolsys.27.1.305

Coley, P. D. (1983). Herbivory and defensive characteristics of tree species in a lowland tropical forest. Ecological Monographs, 53, 209–234. doi: https://doi.org/10.2307/1942495

Connell, J. H. & Slatyer, R. O. (1977). Mechanisms of succession in natural communities and their role in community stability and organization. American Naturalist, 111, 1119–1144. doi: https://doi.org/10.1086/283241

Crawley, M. J. (1983). Herbivory. The dynamics of animal--plant interactions. Blackwell Scientific Publications.

Davis, M. A. Wrage, K. J. & Reich, P. B. (1998). Competition between tree seedlings and herbaceous vegetation: support for a theory of resource supply and demand. Journal of Ecology, 86, 652–661. doi: https://doi.org/10.1046/j.1365-2745.1998.00087.x

Deb, S., Sarkar, A., Majumdar, K. & Deb, D. (2014). Community structure, biodiversity value and management practices of traditional agroforestry systems in Tripura, north-east India. Journal of Biodiversity Management and Forestry, 3, 1–6, doi: 10.4172/2327-4417.1000129.

Denslow, J. S. (1980). Patterns of plant species diversity during succession under different disturbance regimes. Oecologia, 46, 18–21. doi: https://doi.org/10.1007/BF00346960

De Steven, D. (1991). Experiments on mechanisms of tree establishment in old-field succession: seedling survival and growth. Ecology, 72, 1076–1088. doi: https://doi.org/10.2307/1940607

Egler, F. E. (1954). Vegetation science concepts I. Initial floristic composition, a factor in old-field vegetation development. Vegetatio, 4, 412–417.

Fayolle, A., Ouédraogo, D-Y, Ligot, G., Daïnou, K., Bourland, N., Tekam, P. & Doucet, J-L. Differential performance between two timber species in forest logging gaps and in plantations in Central Africa. Forests, 6, 380–394. doi:10.3390/f6020380

Feeny, P. (1976). Plant apparency and chemical defense. In Biochemical interaction between plants and insects (pp. 1–40). Springer US.

Ganesh, T. & Davidar, P. (2001). Dispersal modes of tree species in the wet forests of southern Western Ghats. Current Science, 80, 394–399.

Gleason, H. A. (1917). The structure and development of the plant association. Bulletin of the Torrey Botanical Club, 44, 463–481. doi: 10.2307/2479596

Goodale, U. M., Berlyn, G. P., Gregoire, T. G., Tennakoon, K. U. & Ashton, M. S. (2014). Differences in survival and growth among tropical rain forest pioneer tree seedlings in relation to canopy openness and herbivory. Biotropica, 46, 183–193. doi: 10.1111/btp.12088

Khan, M. L. & Tripathi, R. S. (1989). Effect of soil moisture, soil texture and light intensity on emergence, survival and growth of seedlings of a few sub-tropical trees. Indian Journal of Forestry, 12, 196–204.

Kaplan, E. L. & Meier, P. (1958). Nonparametric estimation from incomplete observations. Journal of the American Statistical Association, 53, 457–81.

Lawrence, D., Suma, V. & Mogea, J. P. (2005). Change in species composition with repeated shifting cultivation: limited role of soil nutrients. Ecological Applications 15, 1952–1967. doi: https://doi.org/10.1890/04-0841

Letcher, S.G., Chazdon, R.L., Andrade, A.C., Bongers, F., van Breugel, M., Finegan, B., Laurance, S.G., Mesquita, R.C., Martínez-Ramos, M. & Williamson, G.B. (2012). Phylogenetic community structure during succession: evidence from three Neotropical forest sites. Perspectives in Plant Ecology, Evolution and Systematics 14, 79–87. doi: https://doi.org/10.1016/j.ppees.2011.09.005

Lin, F., Comita, L. S., Wang, X., Bai, X., Yuan, Z., Xing, D., & Hao, Z. (2014). The contribution of understory light availability and biotic neighborhood to seedling survival in secondary versus old-growth temperate forest. Plant Ecology, 215, 795–807. doi: 10.1007/s11258-014-0332-0

McCook, L. J. (1994). Understanding ecological community succession: causal models and theories, a review. Vegetatio, 110, 115–147. doi: https://doi.org/10.1007/BF00033394

Pickett, S. T. A. (1989). Space-for-time substitution as an alternative to long-term studies. In Longterm studies in ecology (pp. 110–135). Springer New York.

Pickett, S. T. A., Collins, S. L. & Armesto, J. J. (1987). Models, mechanisms and pathways of succession. The Botanical Review, 53, 335–371. doi: https://doi.org/10.1007/BF02858321

Putz, F. E. & Canham, C. D. (1992). Mechanisms of arrested succession in shrublands: root and shoot competition between shrubs and tree seedlings. Forest Ecology and Management, 49, 267–275. doi: https://doi.org/10.1016/0378-1127(92)90140-5

Qureshi, I. M. (1968). The concept of fast growth in forestry and the place of indigenous fast growing broad-leaved species. Indian Forester, 94, 51–56.

R Development Core Team. (2015). R: A language and environment for statistical computing. R Foundation for Statistical Computing, Vienna, Austria. URL https://www.R-project.org/.

Raman, T. R. S., Rawat, G. S. & Johnsingh, A. J. T. (1998). Recovery of tropical rainforest avifauna in relation to vegetation succession following shifting cultivation in Mizoram, northeast India. Journal of Applied Ecology, 34, 214–231. doi: https://doi.org/10.1046/j.1365-2664.1998.00297.x

Riswan, S., Kenworthy, J. B. & Kartawinata, K. (1985). The estimation of temporal processes in tropical rainforest: a study of primary mixed dipterocarp forest in Indonesia. Journal of Tropical Ecology, 1, 171–182. doi: https://doi.org/10.1017/S0266467400000225

Singh, P., Haridasan, K., Borang, A., Bhatt, B., Limboo, D. & Borah, M. (1996). Baseline survey of biodiversity of high priority biologically rich areas of Arunachal Pradesh. Sub-project-Mouling area. a Report to the State Forest Research Institute, Itanagar, and WWF–India, New Delhi, India.

Schleicher, A., Peppler-Lisbach, C., & Kleyer, M. (2011). Functional traits during succession: is plant community assembly trait-driven? Preslia, 83, 347–370.

Sundriyal, R. C., Sharma, E., Rai, L. K. & Rai, S. C. (1994). Tree structure, regeneration and woody biomass removal in a sub-tropical forest of Mamlay watershed in the Sikkim Himalaya. Vegetatio, 113, 53–63. doi: https://doi.org/10.1007/BF00045463

Tabarelli, M. & Peres, C. A. (2002). Abiotic and vertebrate seed dispersal in the Brazilian Atlantic forest: implications for forest regeneration. Biological Conservation, 106, 165–176. doi: https://doi.org/10.1016/S0006-3207(01)00243-9

Teegalapalli, K. & Datta, A. (2016). Field to a forest: patterns of forest recovery following shifting cultivation in the Eastern Himalaya. Forest Ecology and Management, 364, 173–182. doi: 10.1016/j.foreco.2016.01.006

Teegalapalli, K., Mailappa, A. S., Lyngdoh, N., & Lawrence, D. (2018). Recovery of soil macronutrients following shifting cultivation and ethnopedology of the Adi community in the Eastern Himalaya. Soil Use and Management, 34, 249–257. doi: https://doi.org/10.1111/sum.12420

Therneau, T. (2015). A Package for Survival Analysis in S. version 2.38, http://CRAN.R-project.org/package=survival

Uhl, C. (1987). Factors controlling succession following slash-and-burn agriculture in Amazonia. Journal of Ecology, 75, 377–407. doi: 10.2307/2260425

Van Do, T., Osawa, A. & Thang, N. T. (2010). Recovery process of a mountain forest after shifting cultivation in Northwestern Vietnam. Forest Ecology and Management, 259, 1650–1659. doi: 10.1007/s11056-010-9225-9

Walters, M. B. & Reich, P. B. (1996). Are shade tolerance, survival, and growth linked? Low light and nitrogen effects on hardwood seedlings. Ecology, 77, 841–853. doi: https://doi.org/10.2307/2265505

