## Supporting information for "Factors affecting the survival of tree seedlings following shifting cultivation in the Eastern Himalaya"

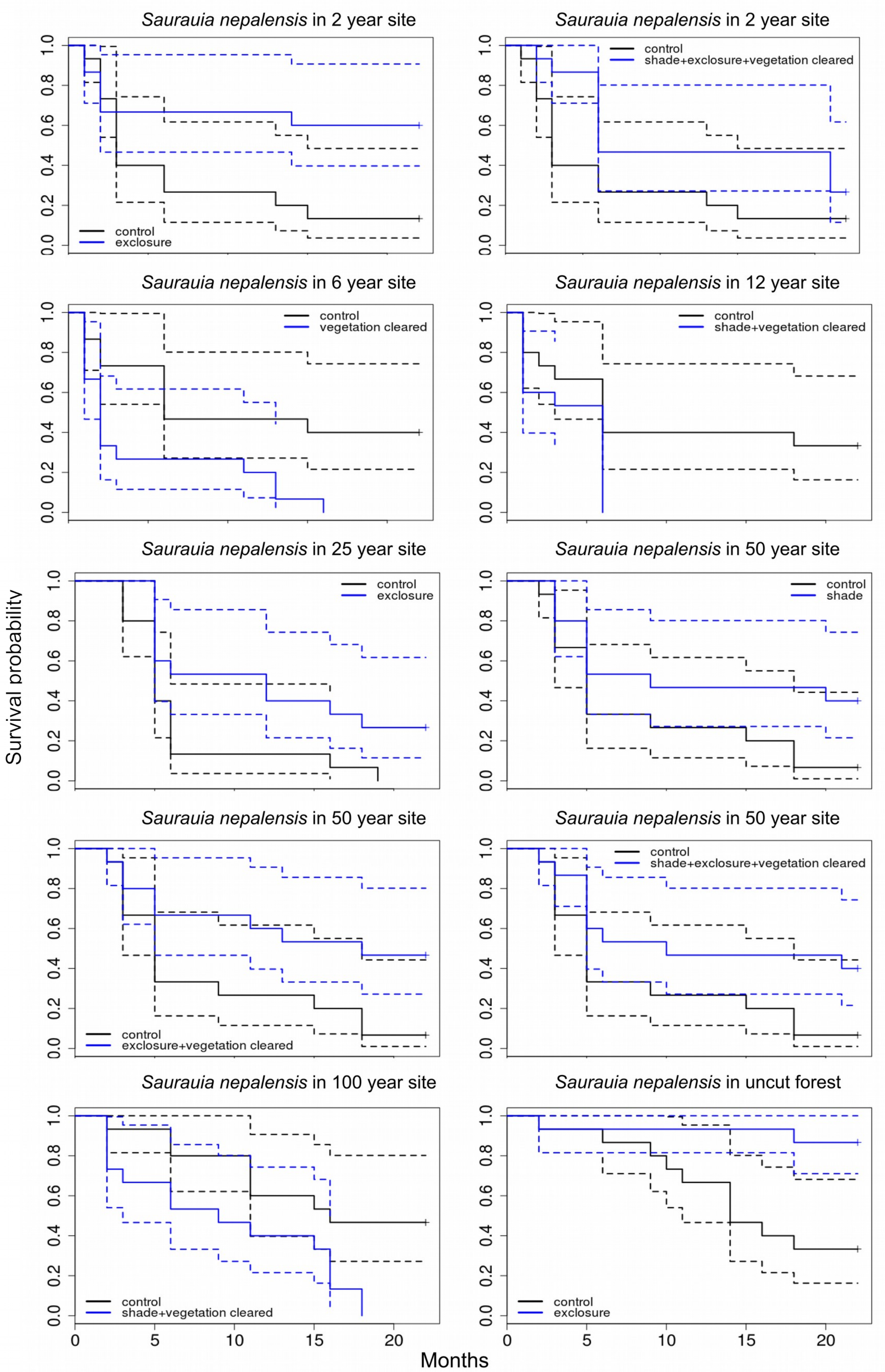


**Figure S1**: Kaplan-Meier seedling survival curves for *Saurauia nepalensis* seedlings for the treatments that were significantly ($p < 0.05$) different from the control plots (dotted lines represent standard errors). The survival probability of seedlings is plotted on the y axis with months on the x axis.

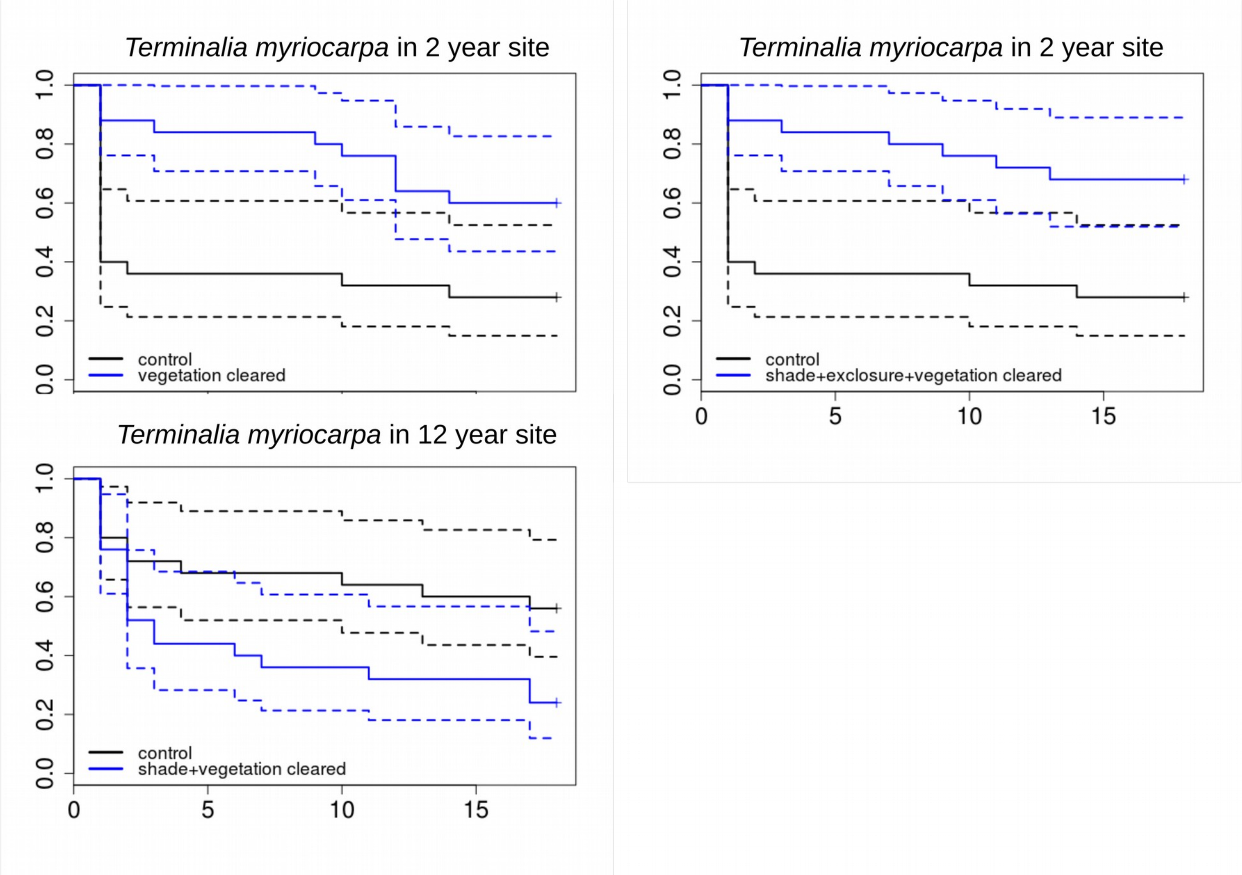


**Figure S2**: Kaplan-Meier seedling survival curves for *Terminalia myriocarpa* seedlings for the treatments that were significantly ($p < 0.05$) different from the control plots (dotted lines represent standard errors). The survival probability of seedlings is plotted on the y axis with months on the x axis.

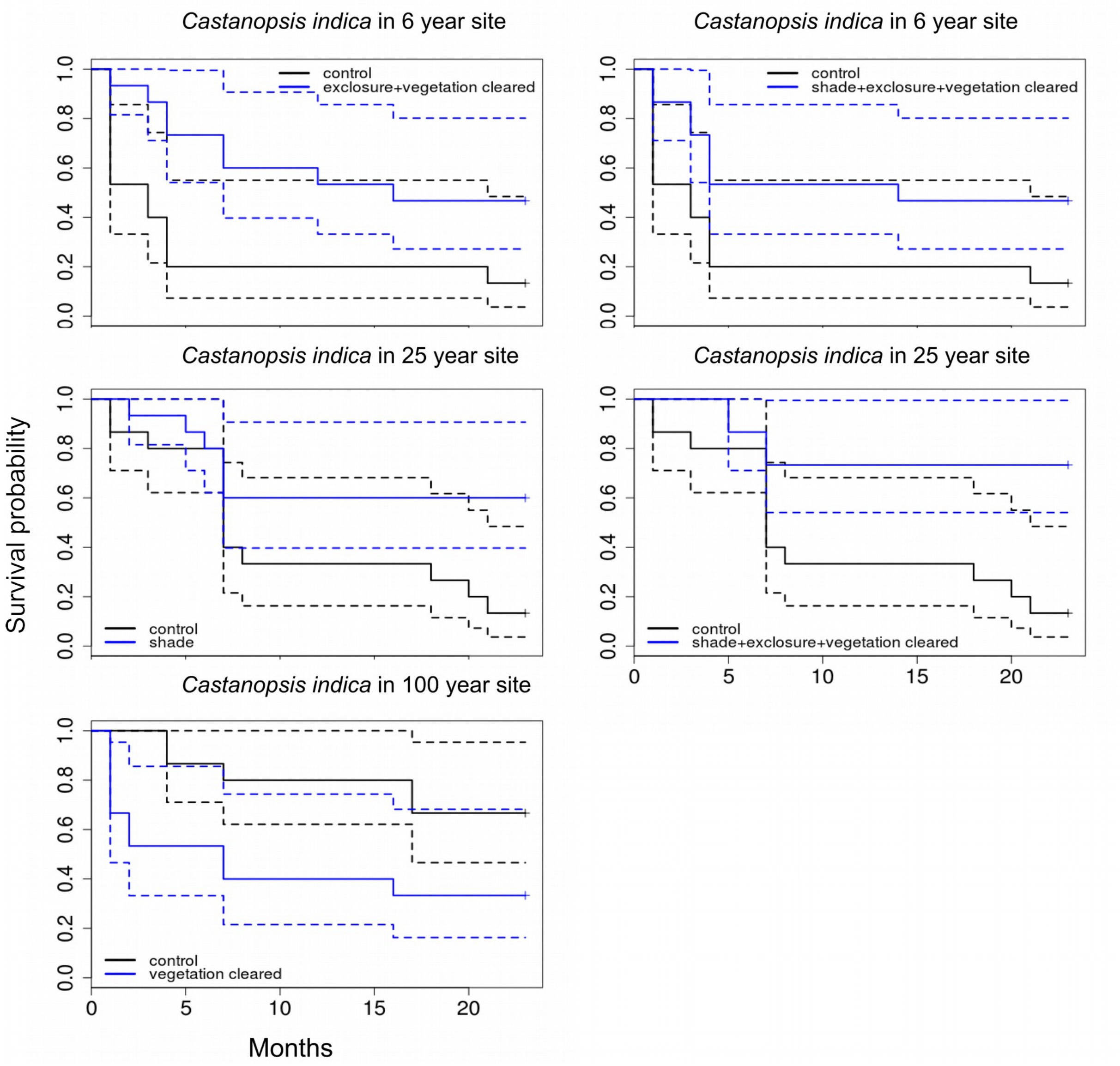


**Figure S3**: Kaplan-Meier seedling survival curves for *Castanopsis indica* seedlings for the treatments that were significantly ($p < 0.05$) different from the control plots (dotted lines represent standard errors). The survival probability of seedlings is plotted on the y axis with months on the x axis.

**Table S1:** Generalised Linear Mixed Models results for seedling survival of *Castanopsis indica* seedlings (pooled across each 1 × 1 m plot) at the end of the study across successional sites formed 2, 6, 12, 25, 50 and 100 years following shifting cultivation and uncut forest and treatments (** indicates a *p*-value < 0.01). Proportion of seedlings alive at the end of the study was the response variable. The factors examined were shade, herbivory and competition with aboveground vegetation. Greenhouse shade netting was used to manipulate available light, exclosures were used to prevent herbivory by mammals and vegetation was cleared to eliminate competition with existing vegetation. Parameter estimates (intercept and contrast), standard errors (SE) and hypothesis tests for parameters are shown.

| ***Castanopsis indica* across all sites and treatments** | | | | |
|---|---|---|---|---|
| | Estimate | S.E. | z value | *p* |
| Age | 0.01 | 0.004 | 2.8 | 0.005** |
| Exclosures | 0.36 | 0.79 | 0.45 | 0.65 |
| Shade | 0.04 | 0.80 | 0.04 | 0.96 |
| Vegetation cleared | -1.48 | 0.91 | -1.62 | 0.10 |
| Shade + Exclosures | -1.03 | 0.86 | -1.20 | 0.23 |
| Shade + Vegetation cleared | -0.29 | 0.81 | -0.35 | 0.72 |
| Exclosures + Vegetation cleared | 0.34 | 0.79 | 0.44 | 0.66 |
| Shade + Exclosures + Vegetation cleared | 1.20 | 0.79 | 1.52 | 0.13 |

**Table S2:** Generalised Linear Mixed Models results for seedling survival of *Saurauia nepalensis* seedlings (pooled across each 1 × 1 m plot) at the end of the study across successional sites formed 2, 6, 12, 25, 50 and 100 years following shifting cultivation and uncut forest and treatments (* indicates a *p*-value < 0.05). Proportion of seedlings alive at the end of the study was the response variable. The factors examined were shade, herbivory and competition with aboveground vegetation. Greenhouse shade netting was used to manipulate available light, exclosures were used to prevent herbivory by mammals and vegetation was cleared to eliminate competition with existing vegetation. Parameter estimates (intercept and contrast), standard errors (SE) and hypothesis tests for parameters are shown.

| ***Saurauia nepalensis* across all sites and treatments** | | | | |
|---|---|---|---|---|
| | Estimate | S.E. | z value | *p* |
| Age | 0.00 | 0.00 | 0.14 | 0.88 |
| Exclosures | 1.81 | 0.72 | 2.50 | 0.01* |
| Shade | 1.19 | 0.70 | 1.68 | 0.09 |
| Vegetation cleared | -5.14 | 5.03 | -1.02 | 0.31 |
| Shade + Exclosures | 0.04 | 0.75 | 0.06 | 0.95 |
| Shade + Vegetation cleared | -0.63 | 0.83 | -0.76 | 0.45 |
| Exclosures + Vegetation cleared | -0.61 | 0.83 | -0.74 | 0.46 |
| Shade + Exclosures + Vegetation cleared | -0.28 | 0.78 | -0.36 | 0.72 |

**Table S3:** Generalised Linear Mixed Models results for seedling survival of *Terminalia myriocarpa* seedlings (pooled across each 1 × 1 m plot) at the end of the study across successional sites formed 2, 6, 12, 25, 50 and 100 years following shifting cultivation and uncut forest and treatments. Proportion of seedlings alive at the end of the study was the response variable. The factors examined were shade, herbivory and competition with aboveground vegetation. Greenhouse shade netting was used to manipulate available light, exclosures were used to prevent herbivory by mammals and vegetation was cleared to eliminate competition with existing vegetation. Parameter estimates (intercept and contrast), standard errors (SE) and hypothesis tests for parameters are shown.

| ***Terminalia myriocarpa* across all sites and treatments** | | | | |
|---|---|---|---|---|
| | Estimate | S.E. | z value | *p* |
| Age | -0.01 | 0.00 | -1.6 | 0.10 |
| Exclosures | -0.23 | 0.68 | -0.33 | 0.74 |
| Shade | -0.49 | 0.70 | -0.70 | 0.48 |
| Vegetation cleared | -0.49 | 0.70 | -0.70 | 0.48 |
| Shade + Exclosures | 0.02 | 0.67 | 0.03 | 0.98 |
| Shade + Vegetation cleared | -0.49 | 0.70 | -0.70 | 0.48 |
| Exclosures + Vegetation cleared | 0.01 | 0.67 | 0.02 | 0.99 |
| Shade + Exclosures + Vegetation cleared | 0.23 | 0.67 | 0.34 | 0.73 |

**Table S4**: Results of the log rank tests to check if the seedling survival curves for control and treatment plots are different for *Saurauia nepalensis* in different-aged sites (age of site in years) and uncut forest (* indicates that the curves are significantly different, when *p*-value ≤ 0.05).

| Age of site | Treatment | N | Chi-sq value | *p*-value |
|---|---|---|---|---|
| 2 | Shade | 15 | 0 | 0.84 |
| 2 | Exclosures | 15 | 4.6 | 0.03* |
| 2 | Vegetation cleared | 15 | 0.1 | 0.75 |
| 2 | Shade + Exclosures | 15 | 3.2 | 0.07 |
| 2 | Exclosures + Vegetation cleared | 15 | 1.7 | 0.20 |
| 2 | Shade + Vegetation cleared | 15 | 1.4 | 0.23 |
| 2 | Shade + Exclosures + Vegetation cleared | 15 | 3.9 | 0.05* |
| 6 | Shade | 15 | 0 | 0.93 |
| 6 | Exclosures | 15 | 3.2 | 0.07 |
| 6 | Vegetation cleared | 15 | 7.9 | 0.00* |
| 6 | Shade + Exclosures | 15 | 1 | 0.33 |
| 6 | Exclosures + Vegetation cleared | 15 | 2.2 | 0.14 |
| 6 | Shade + Vegetation cleared | 15 | 0.3 | 0.56 |
| 6 | Shade + Exclosures + Vegetation cleared | 15 | 0.1 | 0.76 |
| 12 | Shade | 15 | 0.1 | 0.77 |
| 12 | Exclosures | 15 | 0 | 0.89 |
| 12 | Vegetation cleared | 15 | 2.9 | 0.09 |
| 12 | Shade + Exclosures | 15 | 0.4 | 0.54 |
| 12 | Exclosures + Vegetation cleared | 15 | 1.8 | 0.18 |
| 12 | Shade + Vegetation cleared | 15 | 4.3 | 0.04 |
| 12 | Shade + Exclosures + Vegetation cleared | 15 | 0.4 | 0.52 |
| 25 | Shade | 15 | 0.4 | 0.54 |
| 25 | Exclosures | 15 | 5.8 | 0.02* |
| 25 | Vegetation cleared | 15 | 3.6 | 0.06 |
| 25 | Shade + Exclosures | 15 | 0 | 0.83 |
| 25 | Exclosures + Vegetation cleared | 15 | 0.7 | 0.41 |
| 25 | Shade + Vegetation cleared | 15 | 0 | 0.83 |
| 25 | Shade + Exclosures + Vegetation cleared | 15 | 1.4 | 0.24 |
| 50 | Shade | 15 | 4.2 | 0.04 |
| 50 | Exclosures | 15 | 3.7 | 0.05* |
| 50 | Vegetation cleared | 15 | 0.4 | 0.54 |
| 50 | Shade + Exclosures | 15 | 2.5 | 0.11 |
| 50 | Exclosures + Vegetation cleared | 15 | 5.2 | 0.02* |

| | | | | |
|---|---|---|---|---|
| 50 | Shade + Vegetation cleared | 15 | 0.5 | 0.47 |
| 50 | Shade + Exclosures + Vegetation cleared | 15 | 4.5 | 0.03* |
| 100 | Shade | 15 | 0.5 | 0.47 |
| 100 | Exclosures | 15 | 0 | 0.98 |
| 100 | Vegetation cleared | 15 | 2.8 | 0.09 |
| 100 | Shade + Exclosures | 15 | 3.1 | 0.08 |
| 100 | Exclosures + Vegetation cleared | 15 | 1.4 | 0.24 |
| 100 | Shade + Vegetation cleared | 15 | 7.1 | 0.00* |
| 100 | Shade + Exclosures + Vegetation cleared | 15 | 1.4 | 0.24 |
| Uncut | Shade | 15 | 0.2 | 0.68 |
| Uncut | Exclosures | 15 | 8.8 | 0.00* |
| Uncut | Vegetation cleared | 15 | 1.5 | 0.22 |
| Uncut | Shade + Exclosures | 15 | 1 | 0.31 |
| Uncut | Exclosures + Vegetation cleared | 15 | 0.1 | 0.79 |
| Uncut | Shade + Vegetation cleared | 15 | 2.1 | 0.15 |
| Uncut | Shade + Exclosures + Vegetation cleared | 15 | 0 | 0.95 |

**Table S5**: Results of the log rank tests to check if the seedling survival curves for control and treatment plots are different for *Terminalia myriocarpa* in different-aged sites (age of site in years) and uncut forest (* indicates that the curves are significantly different, when p-value ≤ 0.05).

| Age of site | Treatment | N | Chi-sq value | *p*-value |
|---|---|---|---|---|
| 2 | Shade | 25 | 0 | 0.93 |
| 2 | Exclosures | 25 | 0.2 | 0.63 |
| 2 | Vegetation cleared | 25 | 7 | 0.01* |
| 2 | Shade + Exclosures | 25 | 2.1 | 0.14 |
| 2 | Exclosures + Vegetation cleared | 25 | 3.1 | 0.08 |
| 2 | Shade + Vegetation cleared | 25 | 2 | 0.15 |
| 2 | Shade + Exclosures + Vegetation cleared | 25 | 9.2 | 0.00* |
| 6 | Shade | 25 | 0.2 | 0.68 |
| 6 | Exclosures | 25 | 2.5 | 0.12 |
| 6 | Vegetation cleared | 25 | 2.4 | 0.12 |
| 6 | Shade + Exclosures | 25 | 1.4 | 0.23 |
| 6 | Exclosures + Vegetation cleared | 25 | 2.5 | 0.11 |
| 6 | Shade + Vegetation cleared | 25 | 0.2 | 0.67 |
| 6 | Shade + Exclosures + Vegetation cleared | 25 | 1.1 | 0.29 |
| 12 | Shade | 25 | 0.1 | 0.73 |
| 12 | Exclosures | 25 | 0.4 | 0.53 |
| 12 | Vegetation cleared | 25 | 2.7 | 0.10 |
| 12 | Shade + Exclosures | 25 | 0.2 | 0.70 |
| 12 | Exclosures + Vegetation cleared | 25 | 1.7 | 0.19 |
| 12 | Shade + Vegetation cleared | 25 | 4.9 | 0.03* |
| 12 | Shade + Exclosures + Vegetation cleared | 25 | 0 | 0.93 |
| 25 | Shade | 25 | 1.3 | 0.26 |
| 25 | Exclosures | 25 | 0 | 0.97 |
| 25 | Vegetation cleared | 25 | 9.6 | 0.00* |
| 25 | Shade + Exclosures | 25 | 10.6 | 0.00* |
| 25 | Exclosures + Vegetation cleared | 25 | 1 | 0.33 |
| 25 | Shade + Vegetation cleared | 25 | 8.5 | 0.00* |
| 25 | Shade + Exclosures + Vegetation cleared | 25 | 4.5 | 0.03* |
| 50 | Shade | 25 | 0 | 0.94 |
| 50 | Exclosures | 25 | 0.2 | 0.64 |
| 50 | Vegetation cleared | 25 | 0.3 | 0.59 |
| 50 | Shade + Exclosures | 25 | 0 | 0.83 |

| | | | | |
|---|---|---|---|---|
| 50 | Exclosures + Vegetation cleared | 25 | 2.1 | 0.14 |
| 50 | Shade + Vegetation cleared | 25 | 0.9 | 0.33 |
| 50 | Shade + Exclosures + Vegetation cleared | 25 | 6.6 | 0.01* |
| 100 | Shade | 25 | 0.5 | 0.48 |
| 100 | Exclosures | 25 | 0.6 | 0.43 |
| 100 | Vegetation cleared | 25 | 1.6 | 0.20 |
| 100 | Shade + Exclosures | 25 | 1.8 | 0.18 |
| 100 | Exclosures + Vegetation cleared | 25 | 1.8 | 0.18 |
| 100 | Shade + Vegetation cleared | 25 | 13.7 | 0.00* |
| 100 | Shade + Exclosures + Vegetation cleared | 25 | 7.4 | 0.00* |
| Uncut | Shade | 25 | 1.9 | 0.17 |
| Uncut | Exclosures | 25 | 0 | 0.85 |
| Uncut | Vegetation cleared | 25 | 0.3 | 0.58 |
| Uncut | Shade + Exclosures | 25 | 0.5 | 0.49 |
| Uncut | Exclosures + Vegetation cleared | 25 | 1.4 | 0.23 |
| Uncut | Shade + Vegetation cleared | 25 | 2.3 | 0.13 |
| Uncut | Shade + Exclosures + Vegetation cleared | 25 | 1.1 | 0.30 |

**Table S6**: Results of the log rank tests to check if the seedling survival curves for control and treatment plots are different for *Castanopsis indica* in different-aged sites (age of site in years) and uncut forest (* indicates that the curves are significantly different, when p-value ≤ 0.05).

| Age of site | Treatment | N | Chi-sq value | *p*-value |
|---|---|---|---|---|
| 2 | Shade | 15 | 0.2 | 0.64 |
| 2 | Exclosures | 15 | 0.2 | 0.64 |
| 2 | Vegetation cleared | 15 | 1.2 | 0.28 |
| 2 | Shade + Exclosures | 15 | 3.1 | 0.08 |
| 2 | Exclosures + Vegetation cleared | 15 | 0.2 | 0.64 |
| 2 | Shade + Vegetation cleared | 15 | 0.2 | 0.64 |
| 2 | Shade + Exclosures + Vegetation cleared | 15 | 0.2 | 0.64 |
| 6 | Shade | 15 | 0.3 | 0.59 |
| 6 | Exclosures | 15 | 1.8 | 0.18 |
| 6 | Vegetation cleared | 15 | 2.6 | 0.11 |
| 6 | Shade + Exclosures | 15 | 1.2 | 0.27 |
| 6 | Exclosures + Vegetation cleared | 15 | 6.6 | 0.01* |
| 6 | Shade + Vegetation cleared | 15 | 0.5 | 0.50 |
| 6 | Shade + Exclosures + Vegetation cleared | 15 | 4.9 | 0.03* |
| 12 | Shade | 15 | 0.2 | 0.69 |
| 12 | Exclosures | 15 | 1.4 | 0.24 |
| 12 | Vegetation cleared | 15 | 0.1 | 0.74 |
| 12 | Shade + Exclosures | 15 | 1.5 | 0.23 |
| 12 | Exclosures + Vegetation cleared | 15 | 0.2 | 0.69 |
| 12 | Shade + Vegetation cleared | 15 | 2.3 | 0.13 |
| 12 | Shade + Exclosures + Vegetation cleared | 15 | 2 | 0.16 |
| 25 | Shade | 15 | 4.8 | 0.03* |
| 25 | Exclosures | 15 | 0.7 | 0.41 |
| 25 | Vegetation cleared | 15 | 0.6 | 0.42 |
| 25 | Shade + Exclosures | 15 | 2.7 | 0.10 |
| 25 | Exclosures + Vegetation cleared | 15 | 0.6 | 0.44 |
| 25 | Shade + Vegetation cleared | 15 | 0.3 | 0.60 |
| 25 | Shade + Exclosures + Vegetation cleared | 15 | 9.3 | 0.00* |
| 50 | Shade | 15 | 0.2 | 0.67 |
| 50 | Exclosures | 15 | 0.6 | 0.43 |
| 50 | Vegetation cleared | 15 | 0.1 | 0.82 |
| 50 | Shade + Exclosures | 15 | 0.2 | 0.64 |

| | | | | |
|---|---|---|---|---|
| 50 | Exclosures + Vegetation cleared | 15 | 0.1 | 0.82 |
| 50 | Shade + Vegetation cleared | 15 | 0.3 | 0.61 |
| 50 | Shade + Exclosures + Vegetation cleared | 15 | 0 | 0.96 |
| 100 | Shade | 15 | 0.4 | 0.54 |
| 100 | Exclosures | 15 | 0 | 0.99 |
| 100 | Vegetation cleared | 15 | 4.8 | 0.03* |
| 100 | Shade + Exclosures | 15 | 3.8 | 0.05* |
| 100 | Exclosures + Vegetation cleared | 15 | 1.1 | 0.29 |
| 100 | Shade + Vegetation cleared | 15 | 0 | 0.9 |
| 100 | Shade + Exclosures + Vegetation cleared | 15 | 1.5 | 0.22 |
| Uncut | Shade | 15 | 0.8 | 0.37 |
| Uncut | Exclosures | 15 | 1.6 | 0.21 |
| Uncut | Vegetation cleared | 15 | 0.4 | 0.54 |
| Uncut | Shade + Exclosures | 15 | 1.5 | 0.22 |
| Uncut | Exclosures + Vegetation cleared | 15 | 0 | 0.85 |
| Uncut | Shade + Vegetation cleared | 15 | 0 | 0.88 |
| Uncut | Shade + Exclosures + Vegetation cleared | 15 | 0.1 | 0.70 |
